# Microscopic and ultramicroscopic anatomical characteristics of root nodules in *Podocarpus macrophyllus* during development

**DOI:** 10.1101/2022.01.11.475828

**Authors:** Li-Qiong Zhu, Hui-Xin Chen, Li-Jun Zhao, Wei-Xin Jiang

## Abstract

To understand the morphological and structural characteristics of root nodules in *Podocarpus macrophyllus* and their development, this study prepared *P. macrophyllus* root nodule samples at the young, mature, and senescent stages. Optical microscopy and transmission electron microscopy (SEM) revealed that new nodules can be formed on roots and senescent nodules; new nodules formed on the roots are nearly spherical and have an internal structure similar to finite nodules; new nodules on senescent nodules are formed by extension and differentiation of the vascular cylinder of the original nodules; and these new nodules are nested at the base of the original nodules, which create growth space for new nodules by dissociating the cortical tissue; clusters of nodules are formed after extensive accumulation, and the growth pattern is similar to that of infinite nodules; the symbiotic bacteria of *P. macrophyllus* root nodules mainly invade from the epidermal intercellular space of the roots and migrate along the intercellular space of the nodule cortex; infected nodule cortex cells have a well-developed inner membrane system and enlarged and loose nuclei; and unique Frankia vesicles, and rhizobia cysts, and bacteriophages can all develop. Compared with common leguminous and nonleguminous plant nodules, *P. macrophyllus* root nodules are more complex in morphology, structure and composition. From the perspective of plant system evolution, the rhizobium nodules in leguminous angiosperms and Frankia nodules in nonleguminous angiosperms are most likely two branches derived from the nodules in gymnosperms, such as *P. macrophyllus*. The conclusions of this study can provide a theoretical basis for the developmental biology of *P. macrophyllus* root nodules and the evolutionary pattern of plant symbionts.

**Highlights:** We discuss from the perspective of cell developmental biology, the rhizobium nodules in leguminous angiosperms and Frankia nodules in nonleguminous angiosperms are most likely two branches derived from the nodules in gymnosperms, such as *Podocarpus macrophyllus*.

## Introduction

Only leguminous plants and a small number of other plants can form nodules, accounting for only a small proportion of plant species (Roy *et al*., 2020). However, the nitrogen fixation in root nodules provides an enormous contribution to nature by increasing plant yields (Suárez *et al*., 2008; Roy *et al*., 2020), maintaining the ecological balance of the Earth (Cai *et al*., 2018), and reducing pollution (Rejili *et al*., 2012), among other contributions; therefore, researchers are interested in this topic, and related research mainly focuses on the principle of nitrogen fixation in root nodules (Yang *et al*., 2015; Chen *et al*., 2019), nodulation mechanisms(Oldroyd, 2013; Dong *et al*., 2020), genetic characteristics (Soyano *et al*., 2014; Liu *et al*., 2019), and morphology and structure (Maunoury *et al*., 2008; Brett *et al*., 2010). In particular, the rhizobium nodules in leguminous plants and Frankia nodules in nonleguminous plants have attracted the most attention because the former is related to grain yield and quality (Herridge *et al*., 2008), while the latter is closely related to the ecological environment. *Podocarpus macrophyllus* belongs to the genus *Podocarpus* of the family Podocarpaceae and is mainly distributed in various provinces south of the Yangtze River in China. *P. macrophyllus* is an evergreen tree with a beautiful shape, strong resistance to stress, and good greening and medicinal value (Wang, 2000). Studies on *P. macrophyllus* have achieved certain results in terms of morphology (Deng, 2018; Zhu *et al*., 2019), physiology (Huang *et al*., 2018), genetic diversity (Wei, 2012), cultivation (Tian, 2007), pharmacology (Sato *et al*., 2008; Wu *et al*., 2012), and symbiotic microorganisms (Huang *et al*., 2007). However, research on *P. macrophyllus* root nodules started late. First, Huang extracted nitrogen-fixing rhizobia from *P. macrophyllus* root nodules, and then Zhao *et al*. (2014) identified actinomycetes and many fungi in the *P. macrophyllus* root nodules, prompting scholars to study the morphological structure, genetic diversity, and nitrogenase activity of *P. macrophyllus* and their correlations with the environment (Huang *et al*., 2007; Zhao *et al*., 2014). However, few studies have explored the developmental biology of *P. macrophyllus* root nodules. This study mainly observed the microscopic composition of *P. macrophyllus* root nodules at different developmental stages, analysed changes in the occurrence and development of such nodules and their infection, including pathogen invasion into nodule cells’ inner membrane system, and examined the morphology and structure of symbiotic bacteria to elucidate the developmental anatomical characteristics of *P. macrophyllus* root nodules; furthermore, the nodule symbiosis system within spermatophyte is discussed, providing a theoretical basis for research on the evolutionary patterns of nodules.

## Materials and methods

### Research materials

The experimental materials were collected from a *P. macrophyllus* plantation in the demonstration forest of Guangxi Academy of Forestry in Nanning, China, with an age of approximately 15 years. Nanning city is located at 107°45′-108°51′E, 22°13′-23°32′N. The landform is typical of low mountains and hills, with an average elevation of 76.5 m. Nanning belongs to the south subtropical monsoon climate and has sufficient sunshine, little frost, and no snow. The annual average temperature is 21.7□. The average annual rainfall is 1298 mm. The dry and wet seasons are distinct; summer is the rainy season, and winter is the dry season.

In October 2018, six well-growing *P. macrophyllus* plants were selected from the demonstration forest. The soil layer near the base of the trunk was accessed, and three normal lateral roots were cut along each of the four directions of east, west, south, and north. Each branch contained 2-4 grade lateral roots, and nodules at different stages of development were observed under enlarged microscope. Two branches in each direction of each plant were stored in FAA solution, and the remaining 1 branch was put into an incubator and taken to the laboratory for further processing.

### Section preparation

The fresh samples were cleaned with tap water. Ten fine roots with nodules were randomly selected and observed directly using a magnifying glass. The roots were cut into approximately 0.5-cm-long samples to prepare temporary slides with distilled water for observation under an optical microscope, and root hairs, nodule locations, nodule morphology, various developmental stages and other characteristics were recorded. The developmental stage of nodules was determined by colour, i.e., light yellow corresponds to the young stage, yellow indicates the mature stage, and yellow-brown denotes the senescent stage.

Preparation and observation of paraffin sections: According to the method of Zhao *et al*. (2014), 10 lateral roots with nodules stored in FAA solution were selected, cleaned and cut into approximately 1-cm samples to prepare sections, and staining was performed using a mixture of 0.05% toluidine blue and 1% sodium borate. The microstructure of the nodules was observed under an optical microscope.

Transmission electron microscopy (TEM) observation: According to the method of Yang *et al*. (2012), six specimens of *P. macrophyllus* root nodules at different developmental stages stored in FAA were collected to prepare sections. The ultrathin sections were observed, photographed, and recorded under TEM.

## Results

### Invasion location and infection pathway of rhizobia on P. macrophyllus

Under an optical microscope, the root tips of *P. macrophyllus* seedlings were usually enlarged, and the epidermis was covered with a hyaline sheath layer (Fig. 1-a), which contained light yellow bacteria. Dense nodules were found in the mature zone and on superior lateral roots. In addition, root hairs were present on the surface of a small number of young roots and nodules (Fig. 1-b), and some root hairs were curved at the tip and contained light yellow bacteria (Fig. 1-c). In the sections of young root nodules, many bacteria were observed in the cortical cells of root nodules, which were closely attached to the cell wall, and many dark blue *Candida*-like threads were also evident in the intercellular space (Fig. 1-d), which are the infection routes used by symbiotic bacteria to invade the cortical cells of nodules.

**Fig.1.**
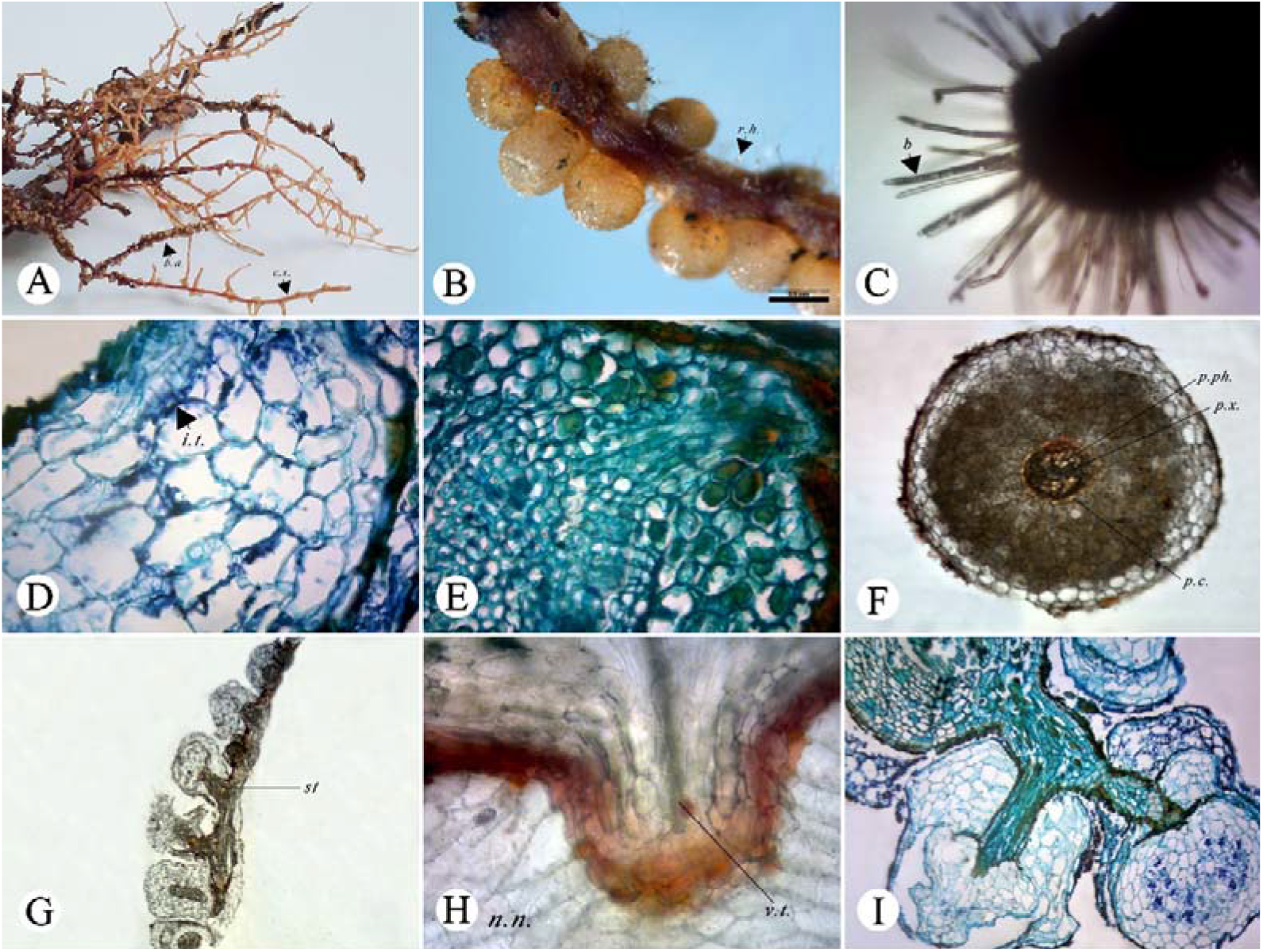
Morphology and microstructure of *Podocarpus macrophyllus* root nodules. (A) Root morphology and arrangement of nodules. b.a-beaded appearance, c.s.-cuticle sheath; (B) The arrow points to the root hair. r.h.-root hair; (C) Morphology of nodule hair (×100), arrow indicates the thallus in nodule hair. b-bacteria; (D) The infection line formed by bacterial invasion of nodule (×400). i.t.-infection thread; (E) A cross section of the root (×400); (F) A cross section of a nodule (×100). p.ph.-primary phloem, p.x.-primary xylem, p.c.-parenchymatous cells; (G) Longitudinal section of roots and nodule (×40), the arrow points to the stele, st-stele; (H) New nodules grow on the old roots (×400) and the vascular tissue of the old root was connected with the vascular tissue of the new nodules. n.n.: new nodule, v.t.:vascular tissue; (I) The stele of the nodule with long branches and nodules with many super imposed layers (×100).

### Occurrence and developmental microanatomy of P. macrophyllus young root nodules

The vascular bundles of young *P. macrophyllus* roots were scattered and consistent with the diarch type. Nodules occurring in the pericycle directly opposite to the primary xylem ridge of the root conformed to the endogenous type, and based on their appearance, the nodules were located on both sides of the root and arranged in two rows (Fig. 1-a, b). According to the appearance of nodules, two types of root nodules can be defined. One type entails root nodules with stalk (Fig. 1-a), i.e., the lateral root primordia and the nodule primordia occur at the same position, the front end of the lateral root is always surrounded by the nodule primordia, and both develop together and break through the root epidermis. The other type refers to root nodules without stalk, where new nodules grow directly from the young roots.

After formation of the cylindrical nodule primordium, periclinal division continues, and nodule primordium breaks through the cortex and epidermis of the young roots. At the same time, nodule primordium divides and differentiates inward to form the vascular tissue of the nodule, which is connected to the vascular tissue of the young root (Fig. 1-e). The vascular tissue is located in the centre of the nodule, conforms to the diarch type, and is composed of xylem, phloem, and parenchyma cells. The xylem tracheids are ring-shaped, and the phloem has sieve cells (Fig. 1-f). The length of the vascular cylinder in the radial section is approximately half of the nodule diameter (Fig. 1-g). The cortical cells of the young roots through which the nodule primordium breaks proliferate, increasing in number and volume, and wrap the vascular tissue of the nodule together to form the nodule cortex. The outermost cells are slightly suberized in the later stage, which confers protection to the nodules. The endothelial layer of root nodules is an extension of the endothelial cells of young roots and forms a closed boundary. The nodules formed on young roots have no meristem or secondary growth and can be classified as determinate nodules.

### Occurrence and developmental microanatomy of P. macrophyllus old root nodules

According to the anatomical diagram (Fig. 1-h), new nodules on old roots originated from the parenchyma inside the root periderm, their dedifferentiation formed nodule primordium, and nodule vascular tissues were formed through periclinal division and connected with the root vascular tissues. Other parenchyma cells around the nodule primordium also divided and expanded to form the nodular cortical parenchyma after stimulation. The innermost parenchyma was densely arranged and became the nodular endothelium, and the nodular outer layer was connected to the root periderm. The cross-sections of old roots and nodules revealed that the nodules still occurred directly opposite to the primary xylem ridge of the roots. The nodules originated near the surface of the roots and belonged to the exogenous type. The nodules had no meristem and secondary growth and can also be classified as determinate nodules.

### Occurrence and developmental microanatomy of P. macrophyllus senescent root nodules

After *P. macrophyllus* root nodules become senescent, some senescent nodules do not detach from the roots, and new nodules can form on them. Longitudinal sections (Fig. 1-i) revealed that cells at the anterior end of the original nodule vascular tissue were dedifferentiated to form a new nodule primordium, its tangential division promoted elongation and growth of the original nodule vascular tissue, and its outward differentiation formed a new nodule cortex. Due to the increase in the volume of the new tissue, the tissue at the tip of the original nodule decomposed and fell off. The base of the new nodule was nested with the remaining part of the original nodule, and the base of the same nodule can have multiple overlaps. Moreover, the dedifferentiation of the original root nodule vascular tissue can occur at two locations on the same cross-section, forming two new nodules. The nodules formed at different levels caused *P. macrophyllus* root nodules to appear clustered. According to the origin and formation of multiple new nodules, these nodules can be classified as indeterminate nodules.

### Ultrastructural characteristics of infected cortical cells and symbiotic microorganisms in P. macrophyllus root nodules at different developmental stages

The cortical cell wall of young root nodules is relatively thick and formed by the superposition of more than 10 layers of band-like structures (Fig. 2-a). Many microorganisms exist in the intercellular space of the cortex (Fig. 2-b), and the morphology of each bacterium is irregular. A cross-section of the bacteria released into the cells was observed (Fig. 2-c), which was approximately circular. Three layers of electron-dense areas were evident on the periphery, with a diameter of approximately 4 μm, and multiple vacuole-like structures, ribosomes, and starch grains were noted inside. Many hyphae were present in the nodules (Fig. 2-d), which differed in structure, including a single-layer structure, a multilayer structure, a branched structure, and a nonbranched structure. Some hyphae are free in the cell, and some are twisted and compressed into a dense mass structure.

**Fig.2.**
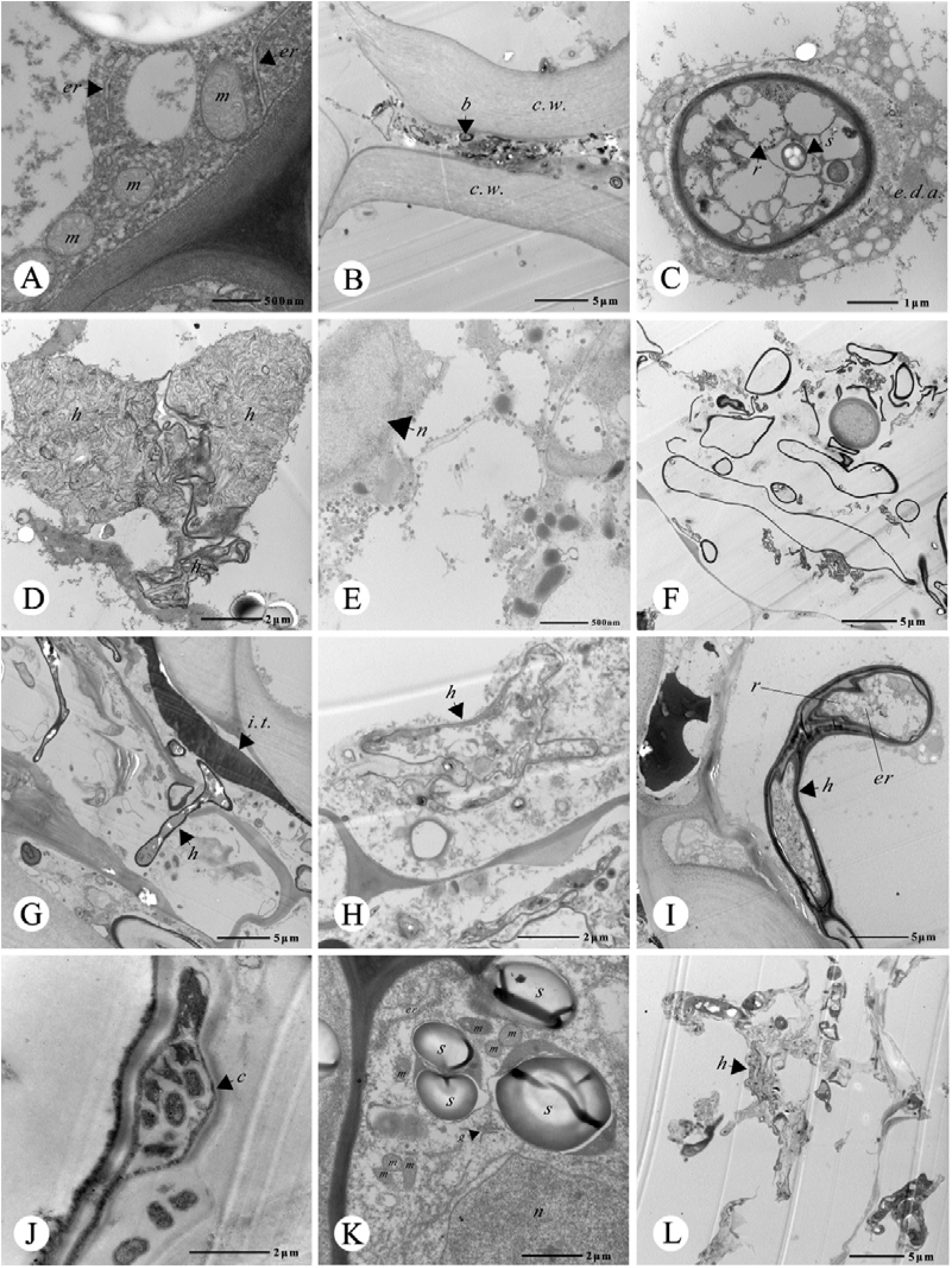
Ultra structure of *Podocarpus macrophyllus* root nodules. (A) The organelles near the cell wall in infected cell of young root nodule(×50000), the arrow points to the endoplasmic reticulum, *er*:endoplasmic reticulum, *m*:mitochondria; (B) Bacteria and multi-layered cell walls in infected cell of young root nodule(×5000), the arrow points to the bacteria, *c*.*w*.: cell wall, *b*: bacteria; (C) Bacteria surrounded by electron dense zone in infected cell of young root nodule(×25000), the arrow points to the ribosome and starch, *r*: ribosome, *s*: starch, *e*.*d*.*a*.: electron dense zone; (D) Dense cluster of arbuscular hyphae in infected cell of young root nodule(×15000), *h*: hyphae; (E) Infected cortical cell of mature root nodule(×100), the arrow points to the nucleus, *n*: nucleus; (F) Hyphae with different form sin infected cell of mature root nodule(×5000); (G) The infection thread in mature root nodule and the process of hyphae invading adjacent cell and releasing bacteria(×5000), the arrow points to the hyphae invading adjacent cell, *i*.*t*.: infection thread, *h*: hyphae; (H) Hyphae in X-shaped(Arrow)(×15000), *h*: hyphae; (I) The top of the hyphae without septum expand stoform a vesicle in infected cell of mature root nodule(×5000), *h*: hyphae, *er*: endoplasmic reticulum, *r*: ribosome; (J) Cyst structure in infected cell of mature root nodule(Arrow)(×20000), *c*: cyst; (K) Starch granules and organelles near the cell wall in infected cell of old root nodule(×15000), the arrow points to the Golgi some, *er*: endoplasmic reticulum, *s*: starch, *n*: nucleus, *m*: mitochondria, *g*: golgiosome; (L) Dissolving hyphae in infected cell of old root nodule(×15000), the arrow points to the hyphae, *h*:hyphae.

Infected cortical cells of mature nodules were rich in organelles, and the nuclei were significantly enlarged and loose (Fig. 2-e). The number of microorganisms was increased, with many diverse forms (Fig. 2-f). The thick, dark, and enlarged linear structure in the intercellular space was the invasion route for symbiotic bacteria. In the cortical cells near the nodules, the hyphae passing through the cell wall could be viewed (Fig. 2-g). The number of intracellular hyphae was increased compared to that in young nodules, with uneven thicknesses and diverse morphologies (Fig. 2-h). Occasionally, vesicles appeared at the front end of the hyphae, and the wall thickness of vesicles was uneven, with 2-3 layers. The vesicles contained organelles such as endoplasmic reticula and ribosomes (Fig. 2-i). A small number of bacteriophage-containing cysts were also found in the cortical cells near the stele (Fig. 2-j). The size of the cyst was approximately 5×2 μm, the diameter of the bacteriophage was 0.5-0.8 μm, and the morphology included ellipsoid, triangular, and irregular shapes.

The number of organelles in the infected cortical cells of senescent nodules was reduced, but nuclei with a loose structure and a small number of mitochondria, Golgi apparatus, endoplasmic reticula, and starch grains were still observed (Fig. 2-k). The colour of endothelial cells was darkened. The hyphae in the cortical cells were loose and irregular in shape. Organelles and inclusions were rare in the cells, vesicles, and cysts, some hyphae had only an empty membrane structure, and the hyphae gradually decomposed in the later period.

## Discussion

### Invasion and migration pathways of symbiotic bacteria

Most plant symbiotic bacteria invade hosts from root hairs, such as in soybean, bayberry, and alder (Zeng, 1987), and some enter from the junction of the root hair base and epidermal cells, such as in lupin (Verma, 1983), while for *Neptunia oleracea*, and *Elaeagnus pungens* (Zeng, 1987), symbiotic bacteria completely enter from the epidermal cells of the host. The proportion of root hairs in *P. macrophyllus* seedlings is very small; therefore, symbiotic bacteria should mainly invade from the epidermal cells of the seedlings. The presence of an infection thread indicates that some of the symbiotic bacteria infect and migrate through the intercellular space, which is consistent with reports by Han *et al*. (1988) and Wang *et al*. (1993). The percentage of infection threads in infected cells has been reported (Mitsustin and Silnikova, 1968) to usually be less than 8%, but the proportion of infection threads observed in this study was higher than this ratio.

### Occurrence and types of nodules

The root nodules formed on young roots originate from the pericycle of the young roots, which share the same origin with the lateral roots. This conclusion is recognized by most researchers (Zeng, 1987). However, this study found that *P. macrophyllus* root nodules had meristem (the meristem is actually a lateral root) (Fig. 1-a), i.e., lateral roots and nodules appear at the same location, and nodules are at located the anterior ends of young roots, indicating that the nodule primordia are located outside the lateral root primordia at the origin. Two possible formation mechanisms can be utilized: (1) After the pericycle is dedifferentiated, the inner cells form the lateral root primordium, the outer cells form the nodule primordium, and both cells then separately develop into the lateral roots and nodules, or one of them. Nutman (1949) agreed with this conclusion. (2) The root nodule primordium is formed by restoring the meristematic capacity of the endothelial parenchyma cells immediately outside the pericycle, and the lateral roots still originate from the pericycle cells. Ren *et al*. (2018) advised that nodules originate from the cortical cells of roots. As mentioned earlier in this paper, new nodules formed on senescent nodules originated from the nodule endothelium at the anterior end of the original nodule vascular tissue (due to extension of the endothelial layer of young roots), and we support the view that nodules on young roots originate from the endothelium of young roots. Xiao *et al*. (2014) also believed that only cells originating from the endothelium of plant roots can be infected by symbiotic bacteria. In terms of structure, the nodules formed on old roots originated from the parenchyma on the inner side of the root periderm. However, we infer that nodules have likely already grown once at this position when the roots are young and fall off due to mechanical friction and other reasons, and the remaining nodules sprout and form new nodules in the later stage. If this is the initiating factor, then the nodules on old and young roots share the same origin.

The above speculation is based only on the developmental anatomical structure of nodules. The next step is to analyse the differential expression of nodules at different locations at different developmental stages to identify the correct initiating factors for nodules.

Root nodules can be divided into two types, i.e., determinate nodules and indeterminate nodules (Kazuhiko, 2011), the former of which have no meristem or secondary growth and are often nearly spherical in shape, and fast-growing root nodules of legumes mostly belong to this type. The latter type has internal meristems and can undergo secondary growth, the nodules often have branching phenomena, and the nodules of alfalfa, pea, clover, bayberry, alder, and other plants belong to this type (Kazuhiko, 2011; Ren, 2018). According to the above classification, the *P. macrophyllus* root nodules formed on young and old roots are finite nodules, while the nodules formed on senescent nodules are infinite nodules. The secondary growth of *P. macrophyllus* root nodules is different from that of common infinite nodules. The meristem of the former belongs to the secondary meristem, and that of the latter belongs to the primary meristem. After entering secondary growth, for the *P. macrophyllus* root nodules, the cortical cells of senescent nodules are damaged and shed to provide growth space for new nodules, resulting in a slightly altered appearance of a single nodule. However, due to the continuous longitudinal extension of the nodule vascular cylinder and the formation of nodule primordia in multiple positions in the horizontal direction, the nodules at some positions appear clustered, which is similar to the appearance of nodules on bayberry, alder, and casuarina (Zeng, 1987), but the principle of formation is different and unique. We infer that if the lifespan of nodules is sufficiently long, the *P. macrophyllus* root nodules formed on young and old roots will very likely enter the secondary growth stage; thus, *P. macrophyllus* root nodules should be classified as indeterminate nodules.

### Changes in various organelles in infected cortical cells at different developmental stages of P. macrophyllus root nodules

In the developmental process of nodules from the young stage to maturity, the inner membrane system of nodule cortex cells increases, and the nuclei become enlarged and loose. Chromosomes can reportedly be transformed into polyploids at this time (Verma, 1983), which was not verified in this study. The number and morphology of mitochondria, endoplasmic reticula, and ribosomes also change with the development of nodules, which is related to the metabolic activities of nodules. Verma *et al*. (1983) and Han *et al*. (1988) showed that with infection by rhizobia, in the cortical cells of nodules, the cytoplasm is gradually thickened, organelles including mitochondria, endoplasmic reticula, and ribosomes continue to increase, and decomposition decreases in the later stage.

### Symbiotic nitrogen-fixing bacteria in P. macrophyllus root nodules

The species, quantity, and morphology of symbiotic bacteria exhibit diversity and uniqueness at each development stage of *P. macrophyllus* root nodules. For a long time, vesicles have been considered a unique marker of Frankia in actinomycetes (Isobel and E. M. S., 1973), and cysts containing bacteriophages are a unique structure of rhizobia (Zeng, 1987). Chen *et al*. (2020) observed hyphae containing vesicles and cysts in *P. macrophyllus* root nodules, and Huang *et al*. isolated rhizobia from *P. macrophyllus* root nodules (Maunoury *et al*., 2008) and confirmed the presence of actinomycetes and rhizobia. The distribution of vesicles and cysts showed that the distribution and nitrogen-fixing areas of the two types of symbiotic bacteria are very likely to be relatively separate, that is, the nitrogen-fixing area of rhizobia is in the middle area of the cortex, and the nitrogen-fixing area of actinomycetes is in the outer area of the cortex, but overlaps may also be present.

From the perspective of morphology and anatomical structure, scattered *P. macrophyllus* root nodules are similar to the fast-growing rhizobium nodules in leguminous plants, and cluster-shaped nodules are similar to the common Frankia nodules in nonleguminous plants, but the *P. macrophyllus* root nodule is unique in terms of the origin, secondary growth, and types of symbiotic bacteria. The combination of genetic sequencing and identification of the lectin contained in *P. macrophyllus* root nodules (Chen, 2020) genetically confirmed that *P. macrophyllus* root nodules have the characteristics of root nodules in both leguminous and nonleguminous plants. In summary, from the perspective of plant symbiosis development, the nodules in both leguminous and nonleguminous angiosperms very likely represent two evolutionary routes derived from nodules in gymnosperms such as *P. macrophyllus*.

## Acknowledgements

We thank professor Ying Hu for providing help during observing the microscopic composition of this research, and professor Bo-Le Jiang for the identification of nitrogen-fixing bacteria. We sincerely thank all those who have provided convenience and help in the process of sampling, writing, translation and submission.

## Author contributions

Li-Qiong Zhu and Li-Jun Zhao designed the research, analyzed the data and wrote the manuscript; Hui-Xin Chen prepared the sample and preformed the experiments.Wei-Xin Jiang revised the manuscript. All authors read and approved the final manuscript.

## References

Brett JF, Arief I, Satomi H, Yu-Hsiang L, Dugald ER, and Peter MG. 2010. Molecular analysis of legume nodule development and autoregulation. Journal of Integrative Plant Biology 52: 61–76.

Cai S, Pittelkow Cameron M, Zhao X, and Wang S. 2018. Winter legume-rice rotations can reduce nitrogen pollution and carbon footprint while maintaining net ecosystem economic benefits. Journal of Cleaner Production 195: 289–300.

Chen HX. 2020. Study on developmental anatony and transcriptomics of Podocarpus macrophyllus root nodules Guangxi University, Nanning, China. (in Chinese)

Chen XP, Lu Q, Liu H, et al. 2019. Sequencing of cultivated peanut, Arachis hypogaea, yields insights into genome evolution and oil improvement. Molecular Plant 12: 920–934.

Deng DL. 2018. Study on anatomical stucture of stems and leaves of male and female plants in three Podocarpus Plants, Guangxi University. (in Chinese)

Dong W, Zhu Y, Chang H, Wang C, Yang J, Shi J, Gao J, Yang W, Lan L, and Wang Y. 2020. An SHR–SCR module specifies legume cortical cell fate to enable nodulation. Nature 589: 586–590.

Han SH. 1988. Ultrastructural study on root nodules of Leguminosae plants. Microbiology China 15: 38–39. (in Chinese)

Herridge DF, Peoples MB, and Boddey RM. 2008. Global inputs of biological nitrogen fixation in agricultural systems. Plant & Soil 311: 1–18.

Huang BL, Lv CQ, Wu B, and Fan LQ. 2007. A Rhizobium isolated from Podocarpus Macrophyllus root nodules. Scientia Sinica(Vitae) 37: 52–57. (in Chinese)

Huang XL, Zhu LQ, Chen HX, He JJ, Huang BY, and Zhao LJ. 2018. Comparative study on seasonal changes in physiological characteristics of three species of Podocarpus. Guangdong Agricultural Sciences 45: 44–49. (in Chinese) 12

Isobel CG, and E. M. S. Gatner. 1973. The formation of vesicles in the developmental cycle of the nodular endophyte of Hippophaë rhamnoides L. Archiv für Mikrobiologie 89: 233–240.

Kazuhiko S. 2011. Rhizobial measures to evade host defense strategies and endogenous threats to persistent symbiotic nitrogen fixation: a focus on two legume-rhizobium model systems. Cellular and Molecular Life Sciences 68: 1327–1339.

Liu CW, Breakspear A, Stacey N, Findlay K, and Murray JD. 2019. A protein complex required for polar growth of rhizobial infection threads. Nature Communications 10: 2848.

Maunoury N, Kondorosi A, Kondorosi E, and Mergaert P. 2008. Cell biology of nodule infection and development, Nitrogen-fixing leguminous symbioses, 153–189. Springer

Mitsustin EN, and Sil’nikova VK. 1968. Biological fixation of atmospheric nitrogen. Macmillan, New York.

Nutman PS. 1949. Physiological studies on nodule formation: II. The Influence of delayed inoculation on the rate of nodulation in red glover. Annals of Botany 13: 261–283.

Oldroyd G. 2013. Speak, friend, and enter: signalling systems that promote beneficial symbiotic associations in plants. Nature Reviews Microbiology 11: 252–263.

Rejili M, Mahdhi M, Fterich A, Dhaoui S, Guefrachi I, Abdeddayem R, and Mars M. 2012. Symbiotic nitrogen fixation of wild legumes in Tunisia: Soil fertility dynamics, field nodulation and nodules effectiveness. Agriculture, Ecosystems and Environment 157: 60–69.

Ren GL. 2018. The evolution of determinate and indeterminate nodules within the Papilionoideae subfamily, Wageningen University

Roy S, Liu W, Nandety RS, Crook A, Mysore KS, Pislariu CI, Frugoli J, Dickstein R, and Udvardi MK. 2020. Celebrating 20 years of genetic discoveries in legume nodulation and symbiotic nitrogen fixation. The Plant Cell 32: 15–41.

Sato K, Sugawara K, Takeuchi H, Park HS, Akiyama T, Koyama T, Aoyagi Y, Takeya K, Tsugane T, and Shimura S. 2008. Antibacterial novel phenolic diterpenes from Podocarpus macrophyllus D. Don. Chemical & Pharmaceutical Bulletin 56: 1691–1697.

Soyano T, Hirakawa H, Sato S, Hayashi M, and Kawaguchi M. 2014. Nodule inception creates a long-distance negative feedback loop involved in homeostatic regulation of nodule organ production. Proc Natl Acad Sci U S A 111: 14607–14612.

Suárez R, Wong A, Ramírez M, Barraza A, M Orozco, Cevallos MA, Lara M, Hernández G, and Iturriaga G. 2008. Improvement of drought tolerance and grain yield in common bean by overexpressing trehalose-6-phosphate synthase in rhizobia. Mol Plant Microbe Interact 21: 958–966.

Tian H. 2007. Seedling-raising technology of Podocarpus Macrophyllus. Foresty and Ecology 5: 23–23. (in Chinese)

Verma DPS. 1983. The molecular biology of Rhizobium-legume symbiosis. International review of cytology. Supplement 14: 211–245.

Wang HY, and Huang WN. 1993. The submicroscopic structure of the endophyte of Elaeagnusconferta Roxb. Acta Phytophysiologica Sinica 19: 61–65. (in Chinese)

Wang JW. 2000. Peculiar trees in Podocarpaceae. Plant Journal 1: 34–36. (in Chinese)

Wei YL. 2012. Study on genetic diversity and rapid propagation of Podocrpus macrophyllus Guangxi University, Nanning, China. (in Chinese)

Wu LD, Liang XH, and Li SY. 2012. Pharmacognostic study on Podocarpus Macrophyllus cinical medical & engineering 19: 52–57. (in Chinese)

Xiao TT, Schilderink S, Moling S, Deinum EE, Kondorosi E, Franssen H, Kulikova O, Niebel A, and Bisseling T. 2014. Fate map of Medicago truncatula root nodules. Development (Cambridge, England) 141: 3517–3528.

Yang K, Tian ZX, Chen CH, et al. 2015. Genome sequencing of adzuki bean (Vigna angularis) provides insight into high starch and low fat accumulation and domestication. Proceedings of the National Academy of Sciences of the United States of America 112: 13213–13218.

Yang YJ. 2012. Biomedical electron microscopy 13-35. The Second Military Medical University Press, Shanghai, China. (in Chinese)

Zeng D. 1987. Nitrogen fixation biology. 151. Xiamen University Press, Xiamen, China. (in Chinese)

Zhao LJ, Zhu LQ, Huang BL, Wen XF, Wei LX, and Sun JL. 2014. Morphology and anatomical structure of root nodules of one-year old Podocarous brevifolius seedlings. Northern Horticulture 10: 53–55. (in Chinese)

Zhu LQ, Deng DL, Zhao LJ, and Li GH. 2019. Comparison on leaf morphological structure of the doecious Podocarpus macrophyllus. Acta Botanica Boreali-Occidentalia Sinica 39: 2179–2186. (in Chinese)

